# Microsporidian obligate intracellular parasites subvert autophagy of infected mammalian host cells to promote their own growth

**DOI:** 10.1101/2022.08.15.503970

**Authors:** Johan Panek, Kacper Sendra, Helen Glenwright, Gregor Kosta, Viktor I. Korolchuk, Robert P. Hirt

## Abstract

Intracellular pathogens such as Microsporidia can interact with host proteostasis pathways such as autophagy. Previous work done in *Caenorhabditis elegans* demonstrated involvement of autophagy in controlling Microsporidian proliferation through ubiquitin labelling of the parasite and subsequent degradation by autolysomes. However, it remains unknown if such mechanisms also play the role in mammalian models. Here we used immunochemistry assays, super-resolution fluorescence microscopy, and modulation of the host’s autophagy flux through siRNA silencing and chemical agents to elucidate how Microsporidia interact with host mammalian cell autophagy flux. We show that despite targeting by early autophagy markers (ubiquitin and p62); the host cell was not able to complete autophagy-mediated removal of *Encephalitozoon cuniculi*. Furthermore, instead of helping to control Microsporidia proliferation, the induction of autophagy dramatically increased proliferation of two Microsporidian species in two different mammalian cell models. Finally, we showed that the reduction of the autophagy flux using siRNA treatment or gut microbiota-derived metabolites known to be important for the intestinal epithelium homeostasis was able to reduce the parasite proliferation. Taken together, our results indicate that Microsporidia infecting mammalian cells not only developed strategies to evade the host autophagy but also evolved way to divert the autophagy flux to promote their own growth.

## Introduction

Protein homeostasis is central to cellular physiology and ultimately to many aspects of health and disease in humans (Taylor et al., 2014). Proteostasis relies on complex cellular regulatory networks that include two major interconnected proteolytic degradation pathways: the ubiquitin-proteasome system and the autophagy-lysosome (autophagy) system (Klaips et al., 2018). Genetic predispositions, ageing, infectious agents, and abiotic environmental factors can all lead to dysregulation of these two pathways, which, in turn, underpins the pathobiology of numerous conditions. This includes infectious (e.g. HIV), neurodegenerative (e.g. Alzheimer’s disease), chronic inflammatory conditions (e.g. inflammatory bowel diseases [IBD] and rheumatoid arthritis), metabolic syndrome-associated diseases and cancers (Balch et al., 2008).

Because they are building their ecological niche inside the host cell, intracellular pathogens interact with host proteostasis, particularly with the autophagy pathway. Indeed, autophagy can play a dual role in the survival of intracellular pathogens by both maintaining nutrient balance of the host that can be exploited by the pathogen to multiply or by directly targeting intracellular pathogens for degradation in a pathway called xenophagy (Sharma et al., 2018). As a result, several of intracellular pathogens developed strategies to either escape this defence mechanism or subvert it. This is best documented for intracellular bacterial pathogens such as *Mycobacterium tuberculosis* or *Listeria spp*. that have developed specific mechanisms/virulent factors to interfere with the maturation of the autophagosome, or *Salmonella spp*. which secretes a deubiquitylation enzyme to remove its tagging by host ubiquitin and thus can evade xenophagy altogether (Bach et al., 2008; Mitchell et al., 2015). In contrast, far less is known about the intracellular eukaryotic parasites and their interaction with host cell autophagy. However, their importance has been increasingly recognized in the recent years, and studies had shown how some microbial eukaryotic parasites have developed strategies to escape and take advantage of autophagy. For example, during its liver stages *Plasmodium spp*. is targeted by host cell xenophagy as attested by rapid labelling of the meronts with host LC3 and SQSTM1/p62 proteins (Schmuckli-Maurer et al., 2017). However, this targeting does not lead to the engulfment of the parasite by canonical autophagosomes. Instead, it is located in the structure that will eventually become the parasitophorous vacuole (PV), suggesting an active recruitment of LC3 and a subversion of xenophagy by the parasite (Wacker et al., 2017). This was confirmed by subsequent studies where induction of autophagy by either starvation or rapamycin treatment in mice increased the parasite load of *Plasmodium berghei* (Prado et al., 2015). Notably, in that same study, genetic suppression of autophagy through an *Atg5* knock out, was both favouring an increase of pathogen load and reducing the size of the parasites, illustrating the dual role of autophagy in controlling the parasite and allowing it to divert nutrients from the host cell for its own benefit (Prado et al., 2015).

Microsporidia is a group of highly diverse strict obligate intracellular pathogens, infecting most animal lineages including economically and ecologically important species (Bateman et al., 2016; Stentiford et al., 2016). Several species can infect humans and represent a significant threat to immunocompromised patients (Mathis et al., 2005). Recent studies have also shown that asymptomatic Microsporidia infections are more common in healthy immune-competent individuals than previously thought (Sak et al., 2011) and may be associated with chronic conditions including Crohn’s disease and cancer (Andreu-Ballester et al., 2013; Leonard et al., 2013). So far, the only study investigating the interaction between Microsporidia and autophagy are available in the *Caenorhabditis elegans* model (Bakowski et al., 2014; Balla et al., 2019). It was suggested that the Microsporidian species *Nematocida parisii* is targeted by *C. elegans* xenophagy machinery in a CUL-6-dependent manner, leading to the labelling of the parasite by host ubiquitin and LC3 (Bakowski et al., 2014). Knockdown of key autophagy genes was sufficient to produce a moderate but significant increase of the parasite load. All these results suggested that in the worm autophagy plays a role in controlling the infection through the xenophagy pathway (Bakowski et al., 2014). In contrast, there is currently no such data in the mammalian systems. In mammals, including humans, intestinal epithelium is the main site of Microsporidia infection (Stentiford et al., 2016). Perturbation of intestinal autophagy, which was shown to be a major cause of intestinal disorder such IBDs or Crohn’s disease, was also linked to Microsporidia infection (Bretin et al., 2016; Hubbard and Cadwell, 2011). Therefore, investigating the interplay between mammalian cell autophagy and the Microsporidian species is of major interest.

We investigated this interplay using *E. cuniculi*, one of the most common Microsporidian species infecting mammals (Katinka et al., 2001). This species was originally isolated from rabbit but is also commonly found in other mammals including humans (Shadduck et al., 1995). The diminutive *E. cuniculi* genome is the results of massive genomic and protein coding capacity reduction (2.2 Mb encoding only 2,000 proteins) leading to the loss of multiple metabolic capabilities (Katinka et al., 2001; Nakjang et al., 2013; Wadi and Reinke, 2020). These include the loss of numerous biosynthesis pathways, and, notably, multiple genes mediating proteostasis, including the mechanistic Target of Rapamycin (mTOR) pathway as well as the majority of the autophagy machinery (Yang et al., 2017). This makes *E. cuniculi* an ideal model system to study the host autophagy response to an eukaryotic intracellular pathogen. Previous transcriptomics analyses found that, similar to *C. elegans*, rabbit kidney cells (RK-13) infected with Microsporidia *T. hominis* (Watson et al., 2015) were characterised by an upregulation of genes tagged as key autophagy components (Bordi et al., 2021). While only *ATG4C* was found to be significantly upregulated in the original dataset (*log*^*2*^ *fold change*:1.59, adjusted *p-value*:0.0025), 10 other genes had increased expression upon infection, including several key *ATG* genes as well as 4 different Cullin genes, suggesting that autophagy/xenophagy may be stimulated upon infection in the mammalian system (Watson et al., 2015). Here we exploited a combination of antibodies developed to investigate *E. cuniculi* infections cycle (Tsaousis et al., 2008), pharmacological treatments and genetic manipulation on mammalian cells to investigate the interplay between Microsporidia infection and mammalian autophagy.

## Results

### *E. cuniculi* co-localizes with host cell ubiquitin and p62 suggesting its targeting by xenophagy machinery

Xenophagy starts with the selective labelling of the pathogens by ubiquitin mediated by an E3 ubiquitin ligase. Ubiquitylation motifs are subsequently recognized by the UBA domain of the receptor protein p62 which, after oligomerization, recruits the autophagy machinery through its LC3-interacting region (LIR) leading to the engulfment of the parasite into an autophagosome (Itakura and Mizushima, 2011). If Microsporidia are targeted by xenophagy in our model, co-localization of the parasite with ubiquitin, p62 and LC3 should be observed. Indeed, previous studies in the *C. elegans* models showed a CUL-6-dependent ubiquitylation of the early stages of the parasite as well as co-localization with the *C. elegans* LC3/ATG8 homolog (LGG-1) (Bakowski et al., 2014).

To verify if such mechanism was conserved in a mammalian model, we investigated co-localization of p62 and conjugated ubiquitin (FK-2) with *E. cuniculi* meronts. The parasite was labelled using an antibody made in a previous study (Tsaousis et al., 2008) which recognizes one of the 4 nucleotide membrane transporters identified in *E. cuniculi* genome (E.c.NTT2). This marker was selected because it is highly expressed during the whole *E. cuniculi* life cycle, from the early meronts to the spore formation stages and also strongly labels the outside of the PV (Fig S1).

Similar to what was seen in the *C. elegans* model, using super resolution fluorescence imaging early meronts were found to be labelled by the conjugated ubiquitin antibody 6 hours post infection (hpi) (Figure 1A-B). Interestingly, only the smaller parasite stages were labelled with ubiquitin (Figure 1C) whereas labelling was never found on later stages suggesting that only early meronts can be targeted by xenophagy. To confirm the targeting of *E. cuniculi* by host autophagy we used the same setup to assay the co-localization of p62 and LC3 6 hpi on the parasite membrane. As expected, parasites were labelled with p62 (Figure 1C-E) however no co-localization with LC3 was found. Therefore, our results suggest that *E. cuniculi* is targeted by host cell xenophagy but developed strategies to escape it by avoiding being isolated into a fully matured autophagic vesicle as evident by the lack of LC3 labelling.

**Figure 1.**
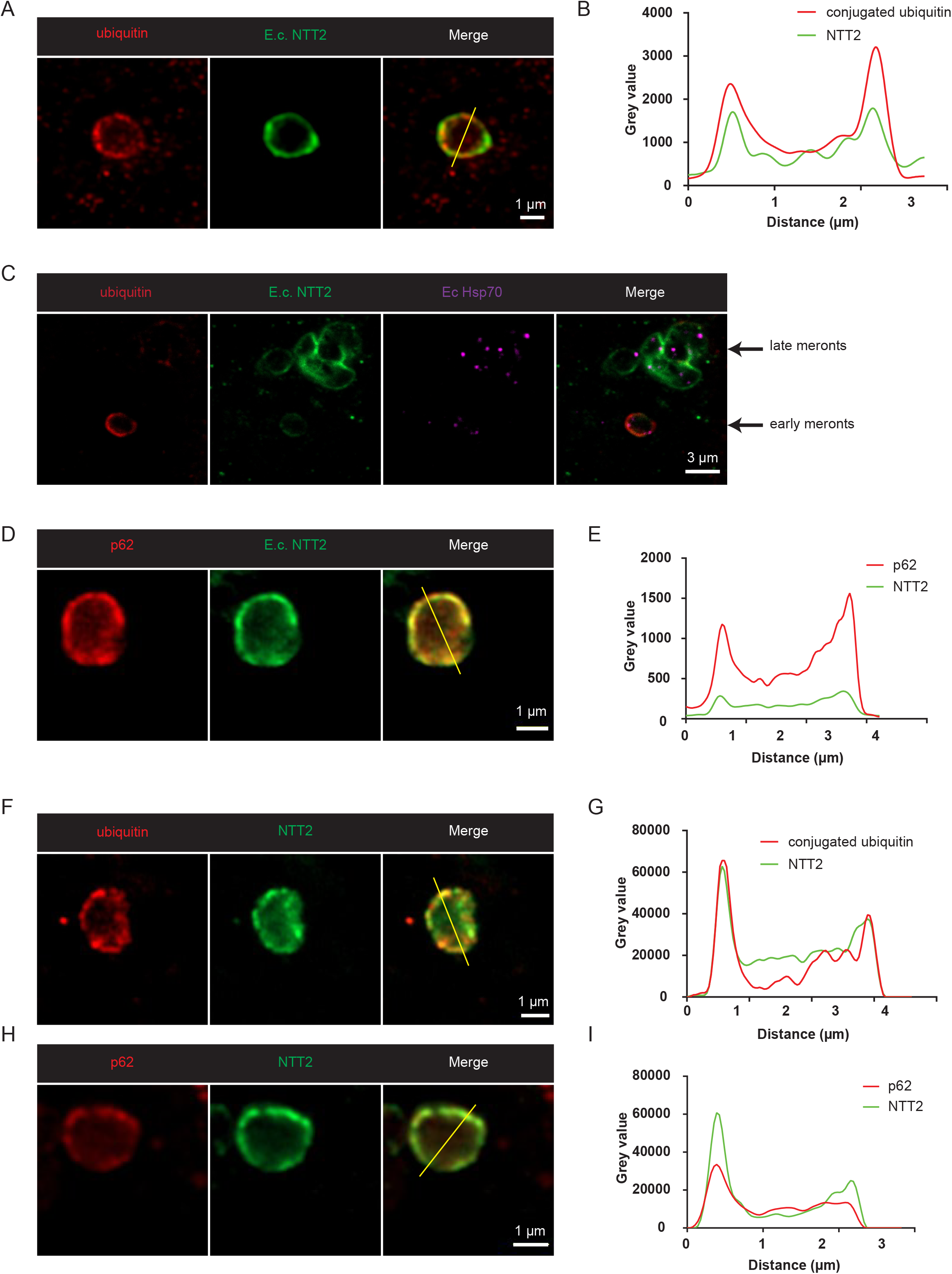
E. cuniculi is targeted by mammalian host cell autophagy. (*A, C, D*) Confocal super resolution imaging (Airyscan) of RK-13 cells 48 hpi by *E. cuniculi*. Parasite PV is labelled in green with α-E.c.NTT2 and in purple with α-HSP70, ubiquitin or p62 are in red. Yellow line indicates transect used for graph B and E. *(B, E)* Plot of the intensity value for green and red signal over a transect indicated in figures A, C and D. (F, H) Confocal super resolution imaging of CACO-2 cells 48 hpi by *E. cuniculi*. Parasite PV is labelled in green with α-E.c.NTT2, ubiquitin or p62 in red. Yellow line indicates transect used for graph G and I. *(G, I)* Plot of the grey value for green and red signal over a transect indicated in figures F and H.

*E. cuniculi* is a natural pathogen of a large range of mammals, where in most cases its infection site is the intestinal epithelium. To confirm our results in a more physiologically-relevant model, we performed the same experiments in human colorectal adenocarcinoma cells (CACO-2 cells), a common model system used to study pathogen-intestinal cell interactions. The results were similar in this cell line with early meronts labelled with ubiquitin, and p62 (Figure 1F-I) while no parasite labelled with LC3.

### Pharmacological induction of autophagy increases *E. cuniculi* proliferation

As xenophagy is not able to clear *E. cuniculi* infection in mammalian cells, we next tested if modulation of the autophagy pathway had any effects on parasite’s infection progression. To assay Microsporidia proliferation, we used two metrics. First, a direct measure of the spore numbers produced per host cell 7 days post infection, and second, the size of the PV 48 hpi. This time point was chosen because the first 2 days represent an initial growth phase during which the parasite increases in size, establishes PV membrane and starts to divide 24 hpi. At 48 hpi new spores start to be produced and released at the centre of the PV, while the newly formed meronts labelled by the E.c.NTT2 antibody form a ring localized to the edge of the PV, allowing for the accurate measurement of its surface (Figure 2A). The PV becomes larger as a direct result of parasite’s cell number increase therefore providing a good proxy for the parasite proliferation. To induce autophagy, we used torin-1, a second-generation mTOR inhibitor which is known to be a very potent autophagy inducer through direct inhibition of both mTORC1 and mTORC2 complexes (Liu et al., 2010). However, due to its high potency, torin-1 has other side-effect such as promoting cell cycle arrest, cells death and protein synthesis arrest (Moschetta et al., 2014). Hence to confirm that the observed effect was due to autophagy induction the other natural mTOR inhibitor rapamycin which only inhibits mTORC1 was also used (Moschetta et al., 2014).

**Figure 2.**
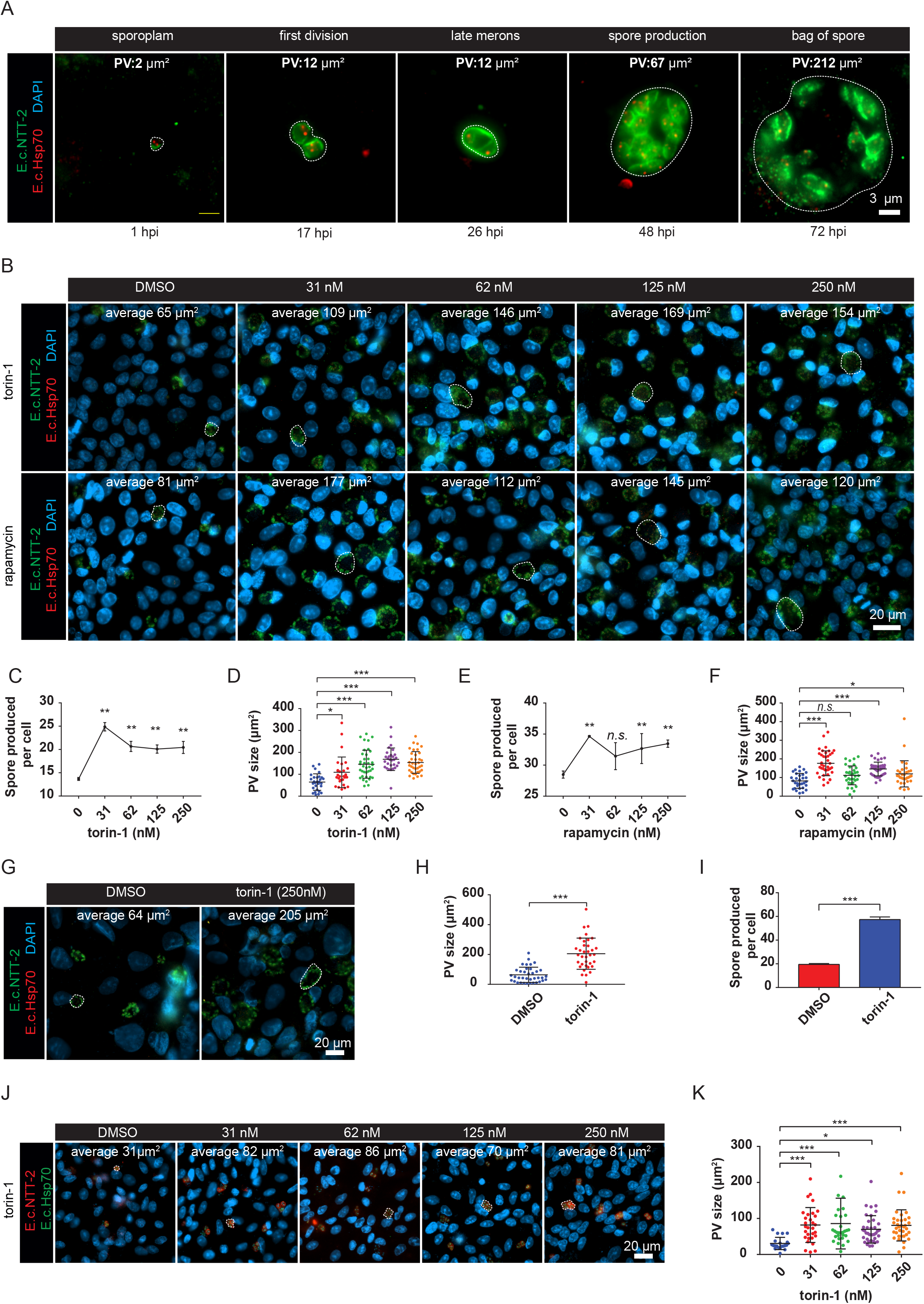
Chemical induction of autophagy increases PV area in RK-13 cells and CAOC-2 cells. *(A)* Fluorescence imaging of RK-13 cell at different time points post infection by *E. cuniculi*, from 6 to 72 hpi. Parasite PV is labelled in green with α-E.c.NTT2 and in red with α-HSP70. PV is highlighted by dashed line and area in µm^2^ measured with ImageJ is indicated on every picture. *(B)* Representative fluorescence imaging of RK-13 cells 48 hpi by *E. cuniculi* in the presence of increasing doses of torin-1 or rapamycin. Parasite PV is labelled in green with α-E.c.NTT2 and in red with α-HSP70. In each picture a representative PV is highlighted with dashed line and the average area of the PV for this condition is indicated. *(C and E)* Graph indicating the number of *E. cuniculi* spores produced per RK-13 cells 7 dpi with different concentrations of torin-1 or rapamycin. 3 independent biological replicates per point, error bars indicate SD. *(D and F)* Graph showing areas of the PV measured in different treatments. Mean area of the replicates is indicated by black bar with errors bars as SD. Each dot represents one measure. At least 30 measurements per condition were done. (G) Representative fluorescence imaging of CACO-2 cells 48 hpi by *E. cuniculi* in presence of 250 nM torin-1 or DMSO. Parasite PV is labelled in green with α-E.c.NTT2 and in red with α-HSP70. In each picture a representative PV is highlighted with dashed line and the average area of the PV for this condition is indicated. *(H)* Graph indicating the number of *E. cuniculi* spores produced per CACO-2 cells 7 dpi with 250 nM torin-1 or DMSO. 3 independent biological replicates per point, error bars indicate SD. *(I)* Areas of the PV measured in the 2 different treatments. Mean area of the replicates is indicated by black bar with error bars as SD. Each dot represents one measure. At least 30 measurements per condition were done. *(J)* Representative fluorescence imaging of RK-13 cells 72 hpi by *T. hominis* in the presence of increasing doses of torin-1. Parasite PV is labelled in red with α-T.h.NTT4 and in green with α-HSP70. In each picture a representative PV is highlighted with dashed line and the average area of the PV for each condition is indicated. *(K)* Graph showing areas of the PV measured in different treatments. Mean area of the replicates is indicated by black bar with errors bars as SD. Each dot represents one measure.

Surprisingly, our results showed that instead of decreasing parasite proliferation both torin-1 and rapamycin increased it. With torin-1 treatment, the spore production per cell 7 dpi is significantly increased, with a maximum reached at the lower tested dose (31 nM) with twice more spores produced than with the DMSO control (Figure 2B-C). The size of the PV is also significantly increased after the treatment with all doses of torin-1 with a maximum reached at 125 nM (Figure 2D). Cells treated with rapamycin showed a similar but milder effect with an increase in spore production (Figure 2B, E) and in size of the PV (Figure 2F) for all but the 62 nM dose. These results suggest that autophagy appears to be beneficial for the parasite proliferation instead of inhibiting it, confirming our previous results indicating that the parasite is not being degraded by xenophagy. The discrepancy between torin-1 and rapamycin treatment effect on parasite growth may be due to the difference in the potency of the drugs to inhibit the mTOR pathway.

Next, we tested whether the ability of the parasite to take advantage of the host autophagy pathway was conserved among different cell lines. Using a similar setup, we tested the effect of the maximum dose of torin-1 (250 nM) on proliferation of *E. cuniculi* in CACO-2 cells. Once again, the results confirmed that torin-1 treatment significantly increases both the number of spores produced per cell and the size of the PV 48 hpi to an even more dramatic extent than in RK-13 cells, with an increase of 300% in spore production compared to control (Figure 2G-I).

To test if other Microsporidian species infecting mammals were able to benefit from the host cell autophagy, we investigated the effect of torin-1 treatment on the growth of *Trachiopleiostophora hominis*, another species of Microsporidia originally isolated from HIV patients (Field et al., 1996)(Heinz et al. 2014). The parasite was labelled with an antibody against T.h.NTT4, one of the 4 NTTs identified in our previous study and which is expressed at high levels during the complete lifecycle of the parasite (Heinz et al. 2014)(Dean et al., 2018). Our results showed a significant increase in area of *the T. hominis* PV upon induction of autophagy by torin-1 treatment similar to the data with *E. cuniculi* in RK-13 and CACO-2 cells (Figure 2J, K).

Taken together these results suggest that instead of controlling the proliferation, autophagy activation in the host cell may benefit both *E. cuniculi* and *T. hominis* proliferation, as attested by the increase in size of the PV and the number of spores produced per cell. This is consistent with our previous results showing that xenophagy targeting of *E. cuniculi* was incomplete (Figure 1). We hypothesized that *E. cuniculi* may have evolved a way to evade xenophagy and, in contrast of being suppressed by it, exploit autophagy to steal nutrients from the host cell. The fact that this phenomenon was observed in two different cell lines and two distantly related parasite species, suggested that this feature might be conserved across different Microsporidia.

### Suppression of autophagy drastically reduces *E. cuniculi* proliferation

Induction of autophagy with torin-1 and rapamycin is indeed a robust and well-characterized approach but may also trigger other adverse effects such cell cycle arrest, protein synthesis arrest and even cell death at higher doses. Hence, to validate our findings we investigated the effect of autophagy inhibition using siRNA targeting a key autophagy gene *ATG5* on *E. cuniculi* proliferation in the presence or absence of torin-1. To take advantage of reagents already developed for human cells, we performed this experiment in CACO-2 cells.

To verify silencing efficiency, ATG5 abundancy, and LC3-II accumulation after Bafilomycin A1 treatment was assayed by immunoblotting in cells treated with either *ATG5* siRNA or non-targeting siRNA (Figure 3A). The results revealed that despite a modest effect on ATG5 levels, the silencing was sufficient to decrease LC3-II accumulation after Bafilomycin A1 treatment by 58% suggesting a strong decrease in the autophagy flux (Figure 3B-C).

**Figure 3.**
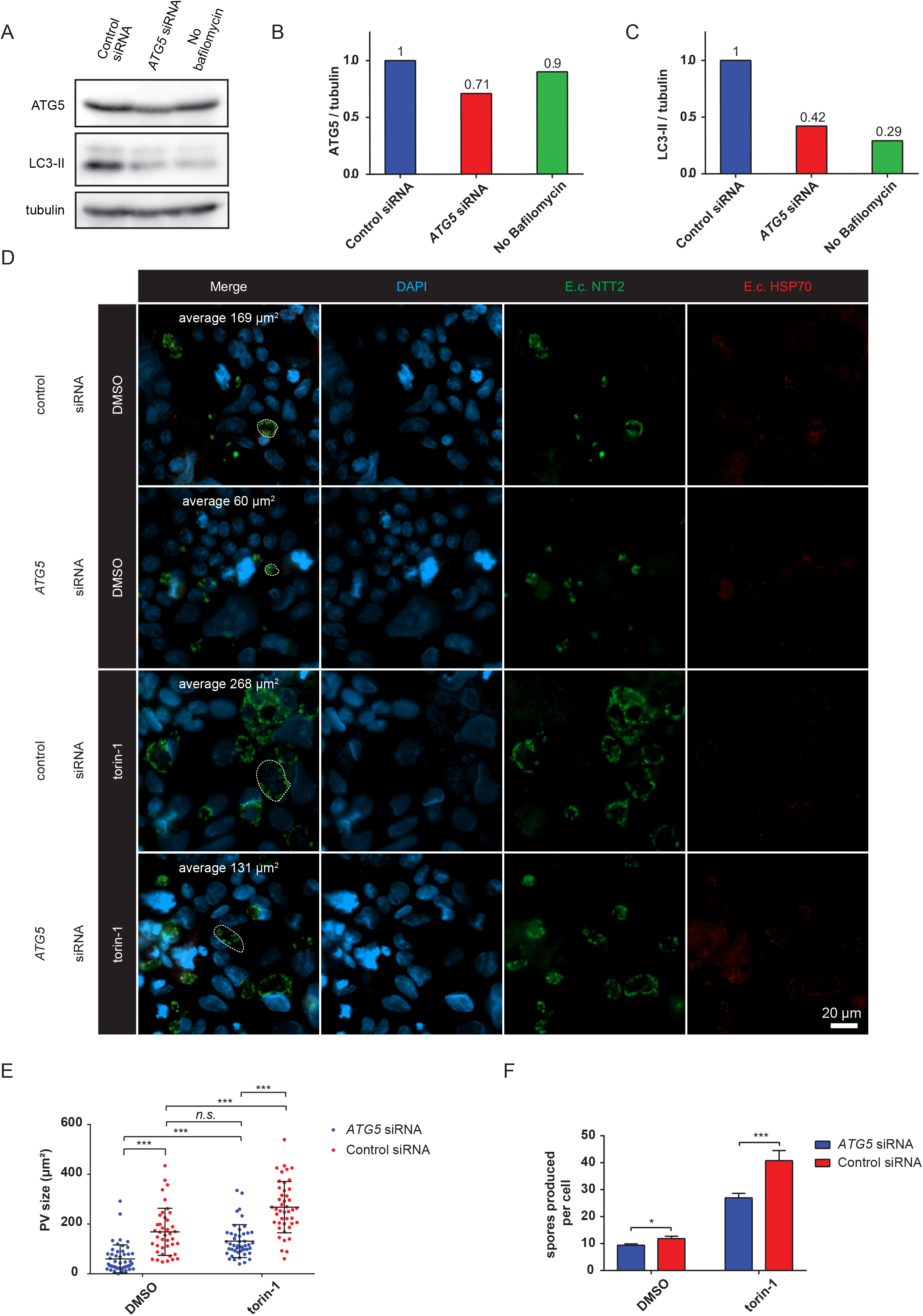
Silencing of *ATG5* by siRNA treatment reduces *E. cuniculi* proliferation in CACO-2 cells. *(A)* Western blot of LC3-II, ATG5 and tubulin (loading control) after ATG5 siRNA treatment of CACO-2 cells. *(B)* Quantification of ATG5 abundance from western blot shown in A. *(C)* Quantification of LC3-II accumulation shown in western blot A. *(D)* Representative fluorescence imaging of RK-13 cells 48 hpi by *E. cuniculi* in the presence in presence or absence of torin-1 (250 nM) and with or without ATG5 siRNA treatment. In each image, a representative PV is highlighted with dashed line and the average area of the PV for this condition is indicated. *(E)* Graph showing areas of the PV measured in the different treatments. Mean area of the replicates is indicated by black bar with errors bars as SD. Each dot represents one measure. *(F)* Graph indicating the number of *E. cuniculi* spores produced per RK-13 cell 7 dpi with different treatments. 3 independent biological replicates per point, error bars indicate SD.

Confocal imaging of CACO-2 cells 48 hpi indicated that *ATG5* silencing significantly reduced the size of *E. cuniculi* PV as well as the number of spores produced per cell in both torin-1 and DMSO treated cells (Figure 3D-F). Furthermore, *ATG5* knockdown reverted the increase of PV area in response to torin-1 treatment (Figure 3D-F). This observation supported the hypothesis that increased PV size and hence parasite proliferation and infection cycle progression after torin-1 treatment are dependent on autophagy induction. The increase in spore numbers in response to torin-1 treatment was also suppressed by *ATG5* knockdown although only partially, which could be explained by these measurements done 7 dpi when the siRNA silencing effect may start to fade away.

### Repression of autophagy with gut microbiota secondary metabolites reduces *E. cuniculi* proliferation

Our results suggest that perturbation of the autophagy flux either due to genetic predisposition such as in Crohn’s Disease (CD), IBD or environmental factors such as microbiota, could have a positive or a negative impact on the susceptibility to Microsporidian infection. Interestingly, a previous study has shown a higher prevalence of Microsporidian infection in CD patients suggesting a potential higher susceptibility to infection, especially to *E. cuniculi* (Andreu-Ballester et al., 2013). Microbiota secondary metabolites can directly affect gut epithelium cell metabolism and physiology. For example, CACO-2 cells exposure to short chain fatty acids (SCFAs), which are naturally produced by the intestinal microbiota in the presence of dietary fibres, has been linked to a decrease in autophagy flux and an increase in the barrier function where both phenotypes could have protective effect against Microsporidian infection (Feng et al., 2018). Hence, we decided to investigate if autophagy flux modulation provoked by incubation with a mix of short chain fatty acids was able to reduce *E. cuniculi* proliferation in CACO-2 cells.

For this experiment, we pre-treated CACO-2 cells with a mix of SCFA (acetate 0.5 mM, butyrate 0.01 mM, and propionate 0.01 mM – as in (Feng et al., 2018)) for 48 h and then infected them with *E. cuniculi*, maintaining the SCFA mix during the entire experiment. Our results confirmed a significant decrease in the size of PV 48 hpi in SCFA treated cells vs control treatment (Figure 4A-B). The number of spores produced per cell also had a trend for reduction in the presence of SCFA, although it did not reach significance (Figure 4C).

**Figure 4.**
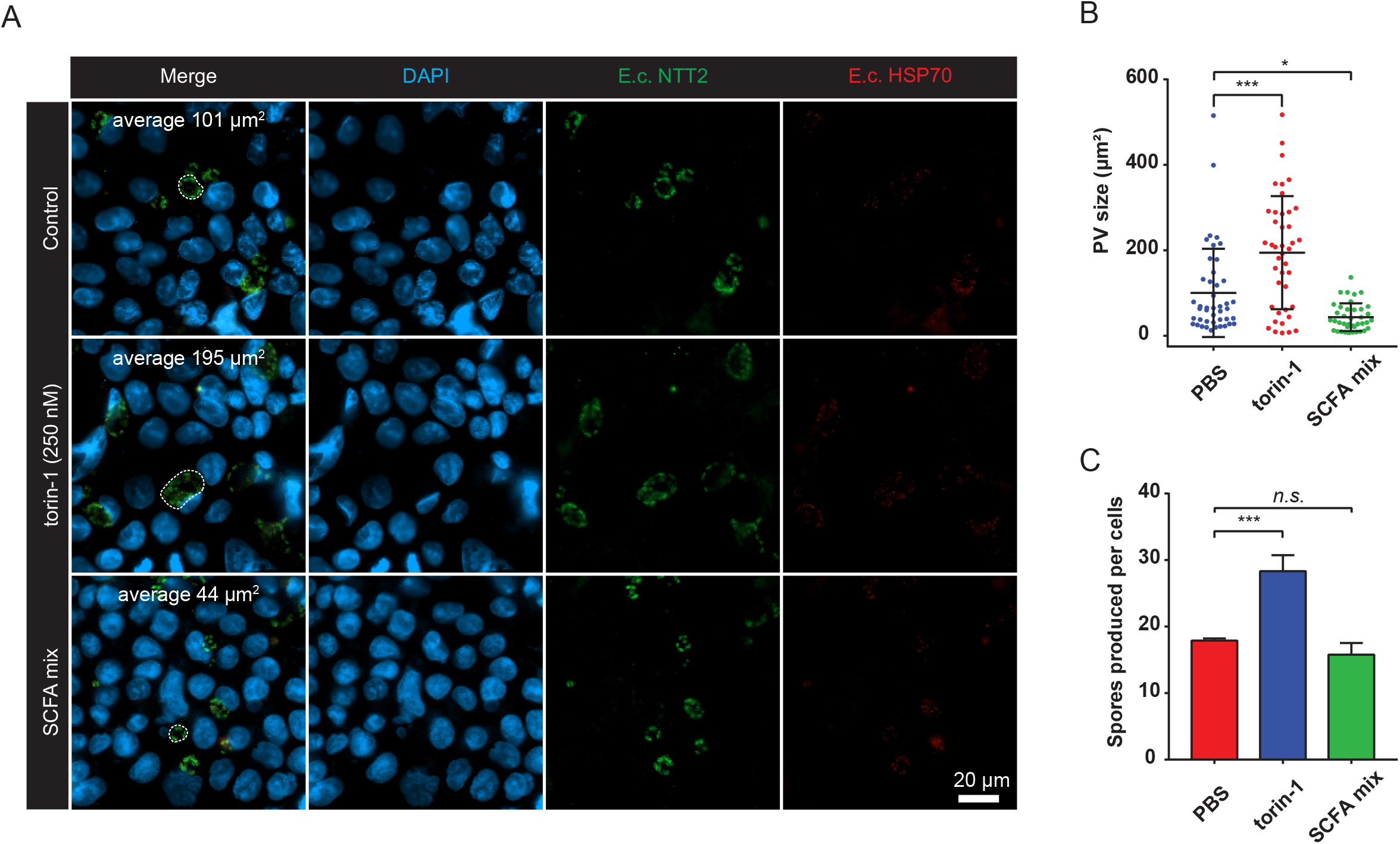
SCFA mix treatment reduces *E. cuniculi* proliferation in RK-13 cells. *(A)* Representative fluorescence imaging of RK-13 cells 48 hpi by *E. cuniculi* in the presence of 250 nM torin-1, SCFA mix or PBS. In each picture, a representative PV is highlighted with dashed line and the average area of the PV for this condition is indicated. *(C)* Graph showing areas of the PV measured in the different treatments. Mean area of the replicates is indicated by black bar with errors bars as SD. Each dot represents one measure. *(D)* Graph indicating the number of *E. cuniculi* spores produced per RK-13 cell 7 dpi with the different treatments. 3 independent biological replicates per point, error bars indicate SD.

## Discussion

Autophagy is a central pathway to respond to cytoplasmic stress, nutrient starvation, and protein aggregation that are all potential outcomes of intracellular pathogen infections. However, to date little has been done to study its involvement in the response of mammalian cells to Microsporidian infection.

Here we found that *E. cuniculi* meronts are partially labelled by the host autophagy markers (ubiquitin and p62) during initial stages of the infection. However, this labelling is not followed by its engulfment into LC3-decorated host cell autophagosome, suggesting that *E. cuniculi* is able to escape xenophagy and avoid degradation. In addition to being apparently protected from xenophagy, the parasite can exploit host autophagy as a source of nutrients as suggested by the significant increase in parasite proliferation following pharmacological induction of autophagy using torin-1 or rapamycin treatment. Notably, our results with another Microsporidia species, *T. hominis*, suggest that this beneficial effect of autophagy induction on Microsporidia proliferation might be conserved in other Microsporidian species infecting mammals. These results were corroborated by our siRNA experiments, where knockdown of *ATG5* reduced the parasite ability to multiply in the human cells. Finally, we showed that microbiota secondary metabolites, namely a mix of three major SCFA, were able to negatively impact *E. cuniculi* multiplication in CACO-2 cells through what we hypothesized to be a modulation of host cell autophagy.

We propose that, unlike in *C. elegans* where autophagy may control Microsporidian infection (Bakowski et al., 2014; Balla et al., 2019), Microsporidia infecting mammalian cells evolved means to escape autophagy and exploit it to support their growth. Indeed, our results clearly indicate that in mammalian cells *E. cuniculi* is able to escape xenophagy despite being labelled by both host ubiquitin and p62, suggesting an evasion mechanism which will need to be elucidated in the future. A similar escape from host xenophagy as well as the beneficial effect of autophagy induction has previously been described for *Plasmodium berghei* infecting mice (Prado et al., 2015). However, in the case of *Plasmodium*, xenophagy suppression (using *Atg5*^-/-^ cell line) was increasing the amount of parasite per cell while reducing their average PV size, suggesting that xenophagy was able to partially control parasite proliferation. In our case, suppression of xenophagy by *ATG5* knockdown in human cells was significantly reducing both the PV size and the parasite proliferation, suggesting that xenophagy might be ineffective in controlling the *E. cuniculi* infection.

Autophagy pathway deregulations are involved in numerous pathologies including IBD(Larabi et al., 2020), a condition that was shown to be associated with Microsporidian infections (Andreu-Ballester et al., 2013). Consistent with these sparse data, a meta-analysis at the IBD-TaMMA platform (Massimino et al., 2021), identified four Microsporidia species, including *E. cuniculi*, to be part of the Fungi significantly more abundant in stools isolated from CD patients compared to healthy controls (Supplementary Table 1). The fact that Microsporidia species were detected in stool samples while they are multiplying in human cells through an unbiased approach (meta-analysis of metatranscriptomics data) further strengthened the association Microsporidia-IBD patients. Several factors could explain this association, for example, patients with established IBD conditions are commonly treated with immunosuppressive drugs that could facilitate Microsporidian infection (Didier and Weiss, 2011). Microsporidia are increasingly appreciated to be ubiquitous in the environment and to infect host from all the animal taxa including humans (Stentiford et al., 2016). Indeed, a recent study identified a high prevalence (14.9%) of infection by the Microsporidian specie *E. bieneusi* among children with IBD treated with various immunosuppressants (Zajączkowska et al., 2021). The altered barrier functions of the epithelium characteristic of IBD (McCole, 2014) could contribute to a higher susceptibility to Microsporidian infections. Here, we show that perturbation of the autophagy flux in intestinal host cells can have direct and dramatic impact on Microsporidian proliferation. On the other hand, by interacting with the host cell autophagy flux and by disturbing the epithelium barrier functions, Microsporidian infections could also represent a potential trigger of the IBDs condition. Hence, Microsporidia could contribute to either, or both, the triggering of IBD and the worsening of its prognosis. Taken together, these different considerations suggest that more interest should be drawn to the potential importance of Microsporidian infections among IBDs patients, which could help at refining IBD patients stratification, a key challenge for these highly heterogenous diseases.

## Methods

### Cell and parasite culture

RK-13 rabbit kidney cell line obtained from the ATCC (ATCC CCL-37) was grown at 37°C with 5% CO_2_ in Dulbecco’s Modified Eagle Medium (DMEM) (Gibco), containing 10% FBS (Gibco), penicillin (100 μg/ml), streptomycin (100 μg/ml), kanamycin (100 μg/ml), and Fungizone (1 μg/ml) purchased from Sigma. CACO-2 cells line obtained from ATCC (ATCC HTB-37) was grown at 37 □C in complete Modified Eagle Medium (MEM) medium complemented with 20% FBS, penicillin (100 μg/ml) streptomycin (100 μg/ml), kanamycin (100 μg/ml) and Fungizone (1 μg/ml). The Microsporidia *E. cuniculi* EC2 and *T. hominis* (ATCC PRA-404) were routinely subcultured in RK-13 cells grown as previously described (Tsaousis et al., 2008). Spores used in time course experiments were harvested from infected RK-13 cells grown in 175 cm^2^ flasks. Cells were washed and scraped in phosphate buffer saline (PBS) and then lysed by 3 passages through a 25G needle and 3 rounds of 1 min sonication on ice. Released spores were purified by layering onto a 25% Percoll gradient and centrifugation at 900 g for 30 min. After 3 washes in PBS, the pelleted spores were resuspended in PBS and used within a week to infect RK-13 or CACO-2 cells.

### Time course experiments

RK-13 cells or CACO-2 cells were seeded into 24-well (for protein extraction) or 6-well plates (for immunofluorescence and spore production assays). When confluent, purified spores were added with a multiplicity of infection of 200. After 30 min the cells were washed 2 times with medium and fresh medium with or without treatment was added (torin-1 31-250 nM *Selleck Chemicals*, rapamycin 12.5-250 nM *Selleck Chemicals*, SCFA mix (acetate 0.5 mM, butyrate 0.01 mM, and propionate 0.01 mM) *Sigma-Aldrich*, DMSO, PBS). Medium was changed every 3 days and treatment was maintained for the full duration of the experiment.

### Spore production assay

7 dpi, cells were washed 2 times with PBS, and detached with trypsin. Part of the cell suspension was used to quantify the number of cells per well. The remaining suspension was centrifuged for 5 min at 16,000 g, the pellet was resuspended in RIPA buffer with 1% SDS. The lysate was sonicated for 1 min at 4 □C to reduce the viscosity and centrifuged for 5 min at 16,000g. The pelleted spores were then resuspended in PBS and counted on Neubauer counting chamber. All assays were done in 3 independent biological triplicates. Spores produced per cell were then calculated and plotted in GraphPad Prism.

### Immunofluorescence assays

6-, 48- or 72-hours post infections, coverslips were washed 3 times with PBS, and fixed for 15 min with ice-cold 50% methanol, 50% acetone. The coverslips were later rehydrated in PBS for 15 min, blocked in PBS 3% milk and labelled with primary antibodies diluted in blocking buffer overnight at 4 □C in humid chamber (mouse α-conjugated ubiquitin - 1:100: *BML-PW8810-0100* - *Enzo life science*, mouse α-p62 - 1:100: *#88588 - Cell Signalling Technology*, rat α-E.c.NTT2 1:200 – [Tsaousis et al., 2008], rabbit α-E.c.HSP70 1:500 – [Tsaousis et al., 2008], rabbit α-T.h.NTT4 1:200 – [Dean et al., 2018], rat α-T.h.HSP70 1:500 – [Dean et al., 2018]). After 3 washes in PBS, coverslips were labelled for 2 h at room temperature with secondary antibodies (α-rat Alexa fluor 488, α-rabbit Alexa fluor 568, α-rabbit Alexa fluor 594 *- Invitrogen*) then washed again 3 times in PBS, counterstained with DAPI and mounted on slides with Prolong glass antifade mounting solution. For co-localisation assays the slides were imaged with a 63X magnification on a Zeiss LSM800 confocal microscope using airyscan super-resolution mode. The red and green signals were then quantified over a transect using ImageJ and intensity values were plotted using GraphPad Prism. For kinetic of infection experiments, the slides were imaged using a Zeiss AxioImager with a 63X magnification. For each condition images were taken randomly on 3 slides using the same acquisition parameters across each set of experiments. The images were then blinded using a home-made python script and the measurement of the parasitophorous vacuoles was done using a home-made ImageJ macro. Briefly, the colour channels were split, the channel containing NTT2 signal was then manually thresholded to separate NTT2 signal from background, the resulting binary image was then used to automatically detect PV using ImageJ detect particles tools. The resulting regions of interest were then measured and saved in a zip file. For each condition and experiment at least 30 PV were measured.

### Immunoblotting

4 h before sampling, Bafilomycin A1 was added in the medium at a final concentration of 400 nM. While on ice, cells were washed twice with ice cold PBS and scrapped on ice in ice-cold RIPA buffer (*Sigma-Aldrich*) with complete Protease Inhibitor Cocktail (*Roche*). After 15 min on ice, cells lysates were centrifuged for 15 min at 16,000g 4 □C and supernatant protein concentration was determined using BCA Protein Assay (*Pierce*). Protein samples were mixed with 4x Laemmli Sample Buffer at a final protein concentration of 5 µg/µl for each sample. 75 µg of proteins were separated on 15% SDS-PAGE gel and transferred on 0.2 µm pore size PVDF membrane. After a blocking step in TBST, 5% milk, membranes were incubated in primary antibodies diluted in TBST, 3% BSA overnight at 4 □C (rabbit α-LC3B 1:1000 – *Cell Signaling/ #4108*, rabbit α-Atg5 1:1000 – *Cell Signaling/#2630*, mouse α-tubulin 1:1000 – *Sigma Aldrich/T5168*). After 3 washes in TBST the membranes were incubated in secondary antibodies for 2 h at room temperature (Goat α-rabbit HRP 1:1000 - *Sigma Aldrich/ GERPN4301*, Goat α-mouse HRP 1:1000– *Sigma Aldrich/GERPN4401*), revealed using ECL reagent (BioRad) and signal was acquired on a BioRad Bioimager. Quantification was done using gel quantification tool in ImageJ.

### siRNA

ON-TARGETplus SMARTpool siRNA against human *ATG5* (L-004374-00-0005) and nontargeting SMARTpool siRNA (D-001810-10-05) were purchased from *Horizon Discovery*. Silencing was done in two steps, a first transfection with a final siRNA concentration of 100 nM was done for 24 h in a 6-well plate, followed by a passaging of the transfected cells into 24-well plate. 24 h after plating, the cells were transfected second time with a final concentration of 20 nM siRNA. 24 h after the second treatment cells were infected and treated with torin-1 (250 nM) or DMSO as a control.

### Statistical analyses

GraphPad Prism version 9.00 was used for statistical analyses and statistical presentation of quantitation. *P-values* are presented in the figures above the compared conditions: *** P < 0.001, ** P < 0.01, * P<0.05. Where two conditions were compared, a paired two-tailed Student’s t test was used (Figures 2H, 2I). If more than two conditions were compared, a one-way ANOVA followed by a Sidak’s multiple comparisons test was applied (Figures 2C, 2D, 2E, 2F, 2K, 4B, 4C). For grouped data, a 2-way ANOVA followed by a Tukey’s multiple comparisons was applied (Figures 3E, 3F).

## Acknowledgements

We are grateful to Newcastle Bioimaging Unit for imaging assistance.

## Author Contributions

J.P., R.P.H., V.I.K. designed research; J.P., K.S., G.K. performed experiments; J.P., K.S. contributed new reagents or analytic tools; J.P. analysed data and wrote the manuscript; R.P.H., V.I.K. edited and proofread the text.

## Figure Legends

**Supplementary Figure 1**. NTT2 transporter is expressed during the whole length of *E. cuniculi* life cycle by meronts localised on the edge of the PV. *(A)* Expression level of NTTs transcript during the *E. cuniculi* time course experiment (data from Grisdale et al., 2013) Km for each of the nucleotide transporter are indicated on the graph (Tsaousis et al., 2008). *(B)* Early meronts labelled with NTT1, NTT2 or NTT4 antibodies. NTT2 antibodies label the periphery of the PV alongside the membrane (white arrowheads). Sporonts (white arrows) labelled with NTT3 antibodies were localized closer to the centre of the vesicle. NTT4 antibodies label small sub-population of cells observed in large PVs corresponding to the late stage of infection but not in any of the time points in the time course experiment. Spores (orange arrowheads) were not labelled with any of the antibodies tested.

**Supplementary Table 1**. Table showing the microbial eukaryote (limited to Fungi) species significantly more abundant (FDR<0.01) in stool-associated tissues from CD patient vs control patient. Microsporidia species are highlighted in red. Data table exported from the IBD Transcriptome and Metatranscriptome Meta-Analysis (IBD-TaMMA) (Massimino et al., 2021) platform that combine data from 26 independent human metatranscriptomics studies with a focus on IBD patients. IBD-TaMMA is accessible through web interface: https://ibd-tamma.readthedocs.io/.

## References

Andreu-Ballester JC, Garcia-Ballesteros C, Amigo V, Ballester F, Gil-Borrás R, Catalán-Serra I, Magnet A, Fenoy S, del Aguila C, Ferrando-Marco J, Cuéllar C. 2013. Microsporidia and its relation to Crohn’s disease. A retrospective study. PLoS One 8:e62107. doi:10.1371/journal.pone.0062107

Bach H, Papavinasasundaram KG, Wong D, Hmama Z, Av-Gay Y. 2008. Mycobacterium tuberculosis Virulence Is Mediated by PtpA Dephosphorylation of Human Vacuolar Protein Sorting 33B. Cell Host Microbe 3:316–322. doi:10.1016/J.CHOM.2008.03.008

Bakowski MA, Desjardins CA, Smelkinson MG, Dunbar TL, Lopez-Moyado IF, Rifkin SA, Cuomo CA, Troemel ER. 2014. Ubiquitin-Mediated Response to Microsporidia and Virus Infection in C. elegans. PLoS Pathog 10:e1004200. doi:10.1371/journal.ppat.1004200

Balch WE, Morimoto RI, Dillin A, Kelly JW. 2008. Adapting Proteostasis for Disease Intervention. Science 319:916–9. doi:10.1126/science.1141448

Balla KM, Lažetić V, Troemel ER. 2019. Natural variation in the roles of C. elegans autophagy components during microsporidia infection. PLoS One 14. doi:10.1371/journal.pone.0216011

Bateman KS, Wiredu-Boakye D, Kerr R, Williams BAP, Stentiford GD. 2016. Single and multi-gene phylogeny of Hepatospora (Microsporidia) - a generalist pathogen of farmed and wild crustacean hosts. Parasitology 1–12. doi:10.1017/S0031182016000433

Bordi M, de Cegli R, Testa B, Nixon RA, Ballabio A, Cecconi F. 2021. A gene toolbox for monitoring autophagy transcription. Cell Death & Disease 2021 12:11 12:1–7. doi:10.1038/s41419-021-04121-9

Bretin A, Carrière J, Dalmasso G, Bergougnoux A, B’chir W, Maurin AC, Müller S, Seibold F, Barnich N, Bruhat A, Darfeuille-Michaud A, Nguyen HTT. 2016. Activation of the EIF2AK4-EIF2A/eIF2α-ATF4 pathway triggers autophagy response to Crohn disease-associated adherent-invasive Escherichia coli infection. Autophagy 12:770–783. doi:10.1080/15548627.2016.1156823

Dean P, Sendra KM, Williams TA, Watson AK, Major P, Nakjang S, Kozhevnikova E, Goldberg A v., Kunji ERS, Hirt RP, Embley TM. 2018. Transporter gene acquisition and innovation in the evolution of Microsporidia intracellular parasites. Nat Commun 9:1–12. doi:10.1038/s41467-018-03923-4

Didier ES, Weiss LM. 2011. Microsporidiosis: Not just in AIDS patients. Curr Opin Infect Dis 24:490–495. doi:10.1097/QCO.0b013e32834aa152

Feng Y, Wang Y, Wang P, Huang Y, Wang F. 2018. Short-Chain Fatty Acids Manifest Stimulative and Protective Effects on Intestinal Barrier Function Through the Inhibition of NLRP3 Inflammasome and Autophagy. Cellular Physiology and Biochemistry 49:190–205. doi:10.1159/000492853

Field AS, Marriott DJ, Milliken ST, Brew BJ, Canning EU, Kench JG, Darveniza P, Harkness JL. 1996. Myositis associated with a newly described microsporidian, Trachipleistophora hominis, in a patient with AIDS. J Clin Microbiol 34:2803–2811.

Grisdale CJ, Bowers LC, Didier ES, Fast NM. 2013. Transcriptome analysis of the parasite Encephalitozoon cuniculi: An in-depth examination of pre-mRNA splicing in a reduced eukaryote. BMC Genomics 14. doi:10.1186/1471-2164-14-207

Hubbard VM, Cadwell K. 2011. Viruses, autophagy genes, and Crohn’s disease. Viruses 3:1281–311. doi:10.3390/v3071281

Itakura E, Mizushima N. 2011. p62 targeting to the autophagosome formation site requires self-oligomerization but not LC3 binding. Journal of Cell Biology 192:17–27. doi:10.1083/jcb.201009067

Katinka MD, Duprat S, Cornillot E, Metenier G, Thomarat F, Prensier G, Barbe V, Peyretaillade E, Brottier P, Wincker P, Delbac F, el Alaoui H, Peyret P, Saurin W, Gouy M, Weissenbach J, Vivares CP. 2001. Genome sequence and gene compaction of the eukaryote parasite Encephalitozoon cuniculi. Nature 414:450–453. doi:http://www.nature.com/nature/journal/v414/n6862/suppinfo/414450a0_S1.html

Klaips CL, Jayaraj GG, Hartl FU. 2018. Pathways of cellular proteostasis in aging and disease. Journal of Cell Biology. doi:10.1083/jcb.201709072

Larabi A, Barnich N, Nguyen HTT. 2020. New insights into the interplay between autophagy, gut microbiota and inflammatory responses in IBD. Autophagy 16:38–51. doi:10.1080/15548627.2019.1635384

Leonard CA, Schell M, Schoborg RV, Hayman JR. 2013. Encephalitozoon intestinalis infection increases host cell mutation frequency. Infect Agent Cancer 8:43. doi:10.1186/1750-9378-8-43

Liu Q, Chang JW, Wang J, Kang SA, Thoreen CC, Markhard A, Hur W, Zhang J, Sim T, Sabatini DM, Gray NS. 2010. Discovery of 1-(4-(4-propionylpiperazin-1-yl)-3-(trifluoromethyl)phenyl)-9-(quinolin-3-yl)benzo[h][1,6]naphthyridin-2(1H)-one as a highly potent, selective Mammalian Target of Rapamycin (mTOR) inhibitor for the treatment of cancer. J Med Chem 53:7146. doi:10.1021/JM101144F

Massimino L, Lamparelli LA, Houshyar Y, D’Alessio S, Peyrin-Biroulet L, Vetrano S, Danese S, Ungaro F. 2021. The Inflammatory Bowel Disease Transcriptome and Metatranscriptome Meta-Analysis (IBD TaMMA) framework. Nature Computational Science 2021 1:8 1:511–515. doi:10.1038/s43588-021-00114-y

Mathis A, Weber R, Deplazes P. 2005. Zoonotic potential of the microsporidia. Clin Microbiol Rev. doi:10.1128/CMR.18.3.423-445.2005

McCole DF. 2014. IBD candidate genes and intestinal barrier regulation. Inflamm Bowel Dis. doi:10.1097/MIB.0000000000000090

Mitchell G, Ge L, Huang Q, Chen C, Kianian S, Roberts MF, Schekman R, Portnoy DA. 2015. Avoidance of Autophagy Mediated by PlcA or ActA Is Required for Listeria monocytogenes Growth in Macrophages. Infect Immun 83:2175. doi:10.1128/IAI.00110-15

Moschetta M, Reale A, Marasco C, Vacca A, Carratù MR. 2014. Therapeutic targeting of the mTOR-signalling pathway in cancer: benefits and limitations. doi:10.1111/bph.12749

Nakjang S, Williams TA, Heinz E, Watson AK, Foster PG, Sendra KM, Heaps SE, Hirt RP, Martin Embley T. 2013. Reduction and expansion in microsporidian genome evolution: new insights from comparative genomics. Genome Biol Evol 5:2285–2303. doi:10.1093/gbe/evt184

Prado M, Eickel N, de Niz M, Heitmann A, Agop-Nersesian C, Wacker R, Schmuckli-Maurer J, Caldelari R, Janse CJ, Khan SM, May J, Meyer CG, Heussler VT. 2015. Long-term live imaging reveals cytosolic immune responses of host hepatocytes against plasmodium infection and parasite escape mechanisms. Autophagy 11:1561–1579. doi:10.1080/15548627.2015.1067361/SUPPL_FILE/KAUP_A_1067361_SM0752.ZIP

Sak B, Kváč M, Kučerová Z, Květoňová D, Saková K. 2011. Latent microsporidial infection in immunocompetent individuals - a longitudinal study. PLoS Negl Trop Dis 5:1–5. doi:10.1371/journal.pntd.0001162

Schmuckli-Maurer J, Reber V, Wacker R, Bindschedler A, Zakher A, Heussler VT. 2017. Inverted recruitment of autophagy proteins to the Plasmodium berghei parasitophorous vacuole membrane. PLoS One 12:e0183797. doi:10.1371/JOURNAL.PONE.0183797

Shadduck JA, Baker MD, Bertucci DC. 1995. Identification and characterization of three Encephalitozoon cuniculi strains. Parasitology 111:411–421. doi:10.1017/S0031182000065914

Sharma V, Verma S, Seranova E, Sarkar S, Kumar D. 2018. Selective Autophagy and Xenophagy in Infection and Disease. Front Cell Dev Biol 6:147. doi:10.3389/FCELL.2018.00147

Stentiford GD, Becnel JJ, Weiss LM, Keeling PJ, Didier ES, Williams BAP, Bjornson S, Kent ML, Freeman MA, Brown MJF, Troemel ER, Roesel K, Sokolova Y, Snowden KF, Solter L. 2016. Microsporidia - Emergent Pathogens in the Global Food Chain. Trends Parasitol 32:1–13. doi:10.1016/j.pt.2015.12.004

Taylor RC, Dillin A, Colby DW, Prusiner SB, Finkbeiner S, Chen B, Retzlaff M, Roos T, Taylor RC, Dillin A, Taylor RC, Dillin A. 2014. Aging as an Event of Proteostasis Collapse 1:1–17. doi:10.1101/cshperspect.a004440

Tsaousis AD, Kunji ERS, Goldberg A v, Lucocq JM, Hirt RP, Embley TM. 2008. A novel route for ATP acquisition by the remnant mitochondria of Encephalitozoon cuniculi. Nature 453:553–556. doi:10.1038/nature06903

Wacker R, Eickel N, Schmuckli-Maurer J, Annoura T, Niklaus L, Khan SM, Guan JL, Heussler VT. 2017. LC3-association with the parasitophorous vacuole membrane of Plasmodium berghei liver stages follows a noncanonical autophagy pathway. Cell Microbiol 19:e12754. doi:10.1111/CMI.12754

Wadi L, Reinke AW. 2020. Evolution of microsporidia: An extremely successful group of eukaryotic intracellular parasites. PLoS Pathog 16:e1008276. doi:10.1371/JOURNAL.PPAT.1008276

Watson AK, Williams TA, Williams BAP, Moore KA, Hirt RP, Embley TM. 2015. Transcriptomic profiling of host-parasite interactions in the microsporidian Trachipleistophora hominis. BMC Genomics 16:1–20. doi:10.1186/s12864-015-1989-z

Yang J, Chai XQ, Zhao XX, Li X. 2017. Comparative genomics revealed the origin and evolution of autophagy pathway. J Syst Evol 55:71–82. doi:10.1111/JSE.12212

Zajączkowska Ż, Akutko K, Kváč M, Sak B, Szydłowicz M, Hendrich AB, Iwańczak B, Kicia M. 2021. Enterocytozoon Bieneusi Infects Children With Inflammatory Bowel Disease Undergoing Immunosuppressive Treatment. Front Med (Lausanne) 8:1717. doi:10.3389/FMED.2021.741751/BIBTEX

