## Supplementary Figure 1 for "Microsporidian obligate intracellular parasites subvert autophagy of infected mammalian host cells to promote their own growth"

Supplemetary Figure 1

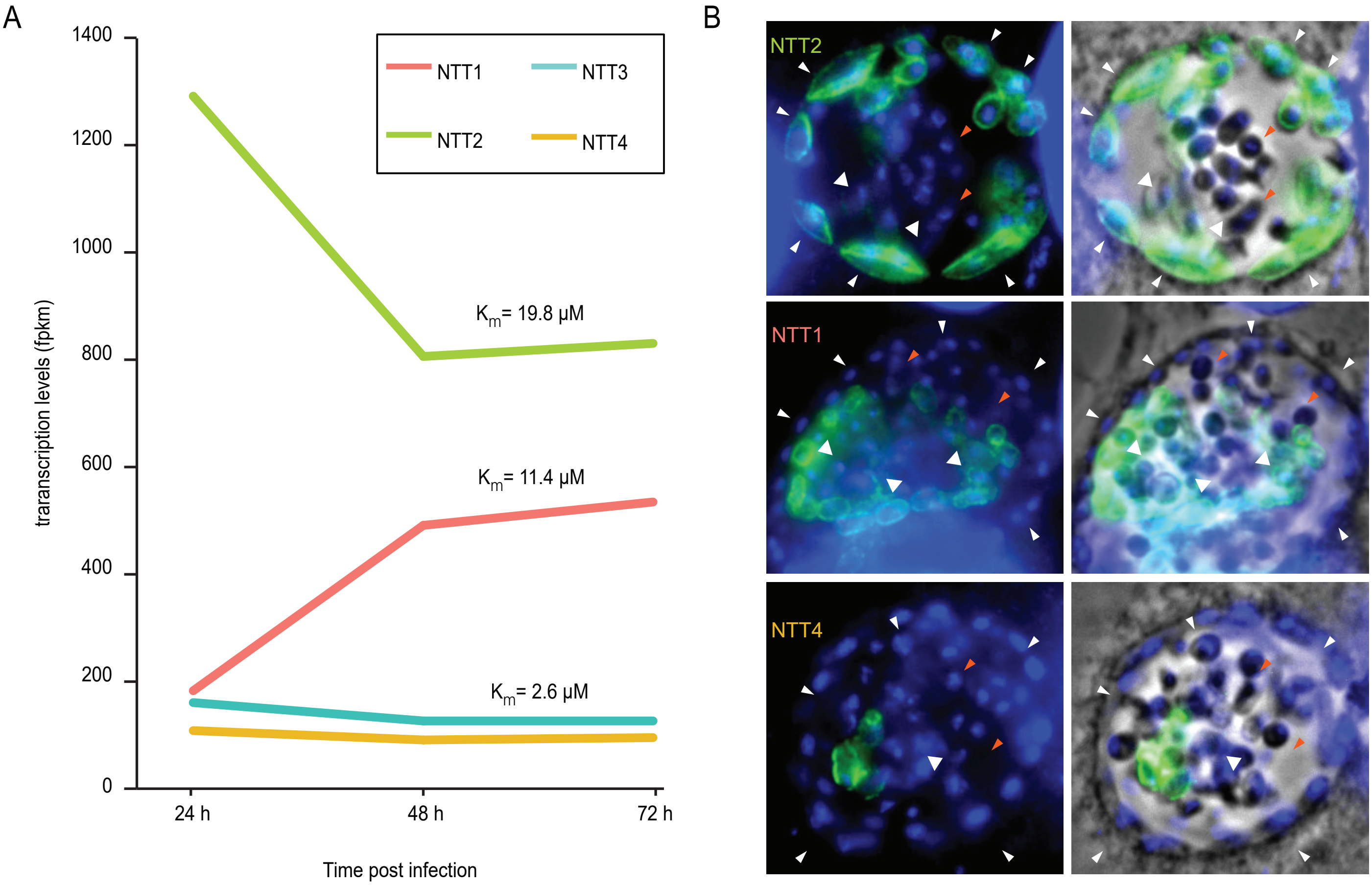


**Supplementary Figure 1.** NTT2 transporter is expressed during the whole length of E. cuniculi life cycle by meronts localised on the edge of the PV. (A) Expression level of NTTs transcript during the E. cuniculi time course experiment (data from Grisdale et al., 2013) Km for each of the nucleotide transporter are indicated on the graph (Tsaousis et al., 2008). (B) Early meronts labelled with NTT1, NTT2 or NTT4 antibodies. NTT2 antibodies label the periphery of the PV alongside the membrane (white arrowheads). Sporonts (white arrows) labelled with NTT3 antibodies were localized closer to the centre of the vesicle. NTT4 antibodies label small sub-population of cells observed in large PVs corresponding to the late stage of infection but not in any of the time points in the time course experiment. Spores (orange arrowheads) were not labelled with any of the antibodies tested.
